# Intronic regulation of SARS-CoV-2 receptor (ACE2) expression mediated by immune signaling and oxidative stress pathways

**DOI:** 10.1101/2021.06.07.447351

**Authors:** Daniel Richard, Pushpanathan Muthuirulan, Jennifer Aguiar, Andrew Doxey, Arinjay Banerjee, Karen Mossman, Jeremy Hirota, Terence D. Capellini

## Abstract

The angiotensin-converting enzyme 2 (ACE2) protein has been highly studied as a key catalytic regulator of the renin-angiotensin system (RAS), involved in fluid homeostasis and blood pressure modulation. In addition to its important physiological role as a broadly-expressed membrane-bound protein, ACE2 serves as a cell-surface receptor for some viruses - most notably, coronaviruses such as SARS-CoV and SARS-CoV-2. Differing levels of ACE2 expression may impact viral susceptibility and subsequent changes to expression may be a pathogenic mechanism of disease risk and manifestation. Therefore, an improved understanding of how *ACE2* expression is regulated at the genomic and transcriptional level may help us understand not only how the effects of pre-existing conditions (e.g., chronic obstructive pulmonary disease) may manifest with increased COVID-19 incidence, but also the mechanisms that regulate ACE2 levels following viral infection. Here, we initially perform bioinformatic analyses of several datasets to generate hypotheses about *ACE2* gene-regulatory mechanisms in the context of immune signaling and chronic oxidative stress. We then identify putative non-coding regulatory elements within *ACE2* intronic regions as potential determinants of *ACE2* expression activity. We perform functional validation of our computational predictions in vitro via targeted CRISPR-Cas9 deletions of the identified *ACE2 cis*-regulatory elements in the context of both immunological stimulation and oxidative stress conditions. We demonstrate that intronic *ACE2* regulatory elements are responsive to both immune signaling and oxidative-stress pathways, and this contributes to our understanding of how expression of this gene may be modulated at both baseline and during immune challenge. Our work supports the further pursuit of these putative mechanisms in our understanding, prevention, and treatment of infection and disease caused by ACE2-utilizing viruses such as SARS-CoV, SARS-CoV-2, and future emerging SARS-related viruses.

**Author Summary:** The recent emergence of the virus SARS-CoV-2 which has caused the COVID-19 pandemic has prompted scientists to intensively study how the virus enters human host cells. This work has revealed a key protein, ACE2, that acts as a receptor permitting the virus to infect cells. Much research has focused on how the virus physically interacts with ACE2, yet little is known on how ACE2 is turned on or off in human cells at the level of the DNA molecule. Understanding this level of regulation may offer additional ways to prevent or lower viral entry into human hosts. Here, we have examined the control of the *ACE2* gene, the DNA sequence that instructs ACE2 protein receptor formation, and we have done so in the context of immune stimulation. We have indeed identified a number of DNA on/off switches for *ACE2* that appear responsive to immuno-logical and oxidative stress. These switches may fine-tune how *ACE2* is turned on or off before, during, and/or after infection by SARS-CoV-2 or other related coronaviruses. Our studies help pave the way for additional functional studies on these switches, and their potential therapeutic targeting in the future.

## Introduction

The angiotensin-converting enzyme 2 (ACE2) protein has been highly studied as a key catalytic regulator of the renin-angiotensin system (RAS), involved in fluid homeostasis and blood pressure modulation(1). ACE2 control on this system occurs both directly (i.e., by lowering levels of angiotensin II) and indirectly (i.e., via alternative cleavage products) inhibiting the self-damaging effects of RAS overactivation, including vasoconstriction, fibrosis, and excessive inflammation(2). The RAS system functions across different organs(1), and similarly, ACE2 is expressed throughout the body(3,4) where it mediates its protective effects and impacts tissue function(2). This activity has prompted the pursuit of ACE2 as a clinical target for protection and treatment against cardiovascular disease, diabetes mellitus, and acute lung damage(2,5).

In addition to its important physiological role as a broadly-expressed membrane-bound protein(2), ACE2 serves as a cell-surface receptor for some viruses - most notably, coronaviruses such as SARS-CoV(6) and SARS-CoV-2(2,5,7). Protein over-expression studies have demonstrated that ACE2 facilitates SARS-CoV-2 infection(7), while mice with engineered human ACE2 are susceptible to infection(8), and it has been suggested that the distribution of ACE2 receptor expression across different tissues contributes to differential virus susceptibility (e.g., lung tissue and alveolar cells)(9). The tissue expression of ACE2 may also explain the wide-ranging symptoms of COVID-19 in patients(10), though alternative means of viral entry have been suggested(4). During infection, ACE2 proteins bound by SARS-CoV-2 particles are endocytosed which, along with increased ADAM17 activity and upstream transcriptional changes, lead to a depletion of cell-surface ACE2 localization and reduced angiotensin catalytic activity(5,10). It has been suggested that reduced expression of *ACE2* may lead to an imbalance of the RAS system in COVID-19 patients, which may represent a major pathological outcome of viral infection(2). These findings have prompted intense interest in the use of recombinant ACE2 and other synthetic mimics as potential therapeutics(11–13).

Given that levels of ACE2 expression may impact viral susceptibility(4,14), and that subsequent changes to expression is a likely pathogenic mechanism of disease(2), an improved understanding of how ACE2 expression is regulated at the genomic and transcriptional level may help us understand not only how the effects of pre-existing conditions (e.g. chronic obstructive pulmonary disease, (COPD)) may manifest with increased COVID-19 incidence, but also the mechanisms that regulate ACE2 levels following viral infection. In this study, we first perform bioinformatic analyses of several datasets to generate hypotheses about *ACE2* gene-regulatory mechanisms in the context of immune signaling and chronic oxidative stress. We next identify putative non-coding regulatory elements within the intronic regions of the *ACE2* gene as potential determinants of *ACE2* expression activity. We then perform functional validation of our computational predictions via targeted deletion of the identified *ACE2 cis*-regulatory elements in the context of immunological stimulation and oxidative stress conditions. Our results demonstrate the presence of intronic *ACE2* regulatory elements responsive to both immune signaling and oxidative-stress pathways, contributing to our understanding of how expression of this gene may be modulated at both baseline and during immune challenge. Furthermore, our work supports the further pursuit of these putative mechanisms in our understanding, prevention, and treatment of infection and disease caused by ACE2-utilizing viruses such as SARS-CoV, SARS-CoV-2, and future emerging SARS-related viruses(15).

## Results

### Up-regulation of *ACE2* gene expression in healthy individuals is associated with immune signalling and viral infection

To first examine patterns of baseline *ACE2* gene expression we analyzed microarray expression datasets from a cohort of healthy, never-smokers (N=109) (see Table S1 for accessions). In these individuals, *ACE2* was co-expressed with a gene set that is most significantly enriched in immune signaling and virus perturbations (Figure 1, Table S1). The top transcription factors associated with these genes included IRFs and STATs (e.g., *IRF1* and *STAT1*). Consistent with this finding, both *IRF1* and *STAT1* genes were also among the top 200 *ACE2* correlated genes. Other genes that were associated with these enriched ‘immune-response’, and ‘viral response’ terms, and co-expressed with *ACE2*, include *IFI16, IFI44, IFI35, NLRC5*, and *TLR3*. These findings suggest that in healthy never-smokers, *ACE2* may be a component of an immune signaling pathway, specifically relating to viral sensing and response and potentially mediated by IRF and STAT transcription factors.

**Figure 1.**
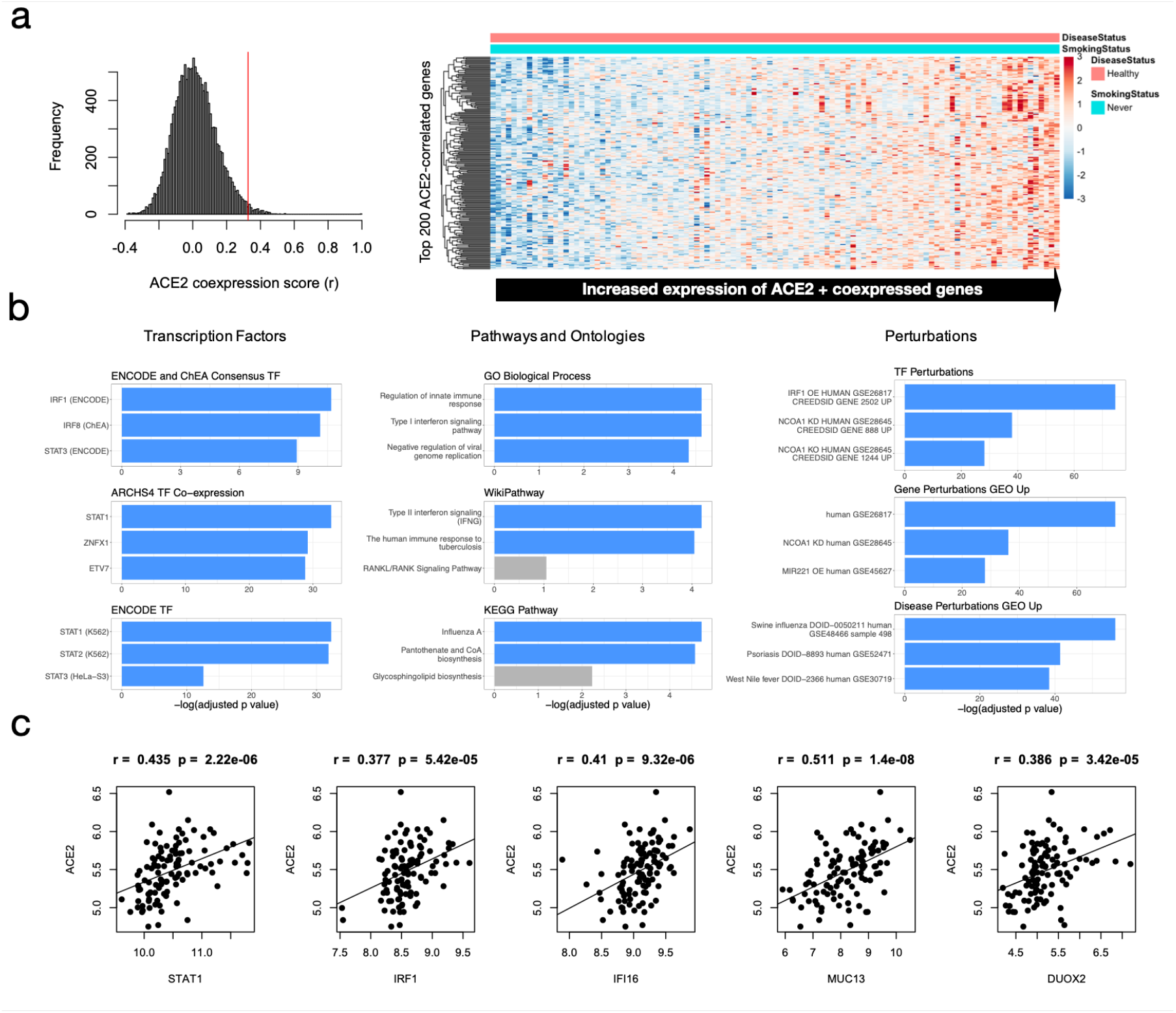
Expression and functional enrichment analysis of *ACE2* and co-expressed genes. **a)** Expression of top 200 *ACE2*-correlated genes (including *ACE2*) in healthy, non-smokers (N=109). **b)** Functional enrichment analysis of top 200 *ACE2*-correlated genes (including *ACE2*). Terms are ranked by -log_2_(FDR-adjusted *p* value) for nine ontologies/groups of interest. **c)** Pearson correlation of *ACE2* with important interferon-related candidate genes found to be co-expressed with *ACE2*.

During our analysis, we also detected as co-expressed with *ACE2, DUOX2*, a known response factor to reactive oxygen species (ROS)(35). This suggests that oxidative stress may be another important mechanism that regulates *ACE2* expression (see also Figure 3). Also of interest are genes that help identify cell-type-specific regulation of *ACE2*, along with its co-regulated gene network. The third top-correlated gene with *ACE2* in healthy non-smokers was *MUC13*, an epithelial mucin known to be expressed in the large intestine and trachea(36) as well as in goblet cells, which are all proposed sites of SARS-CoV-2 replication(2).

We next examined a cohort of N = 136 individuals with asthma(37) to investigate whether the observed associations persisted in individuals with chronic inflammatory lung disease (Figure S1). *ACE2* co-expression in asthmatic individuals was also associated with immune signaling, antiviral responses, and IRF and STAT transcription factors. The top *ACE2*-correlated gene in asthmatics was *CD47*, which is involved in the regulation of interferon gamma. Consistent with this finding, *ACE2* and *CD47* are both co-expressed with the interferon-inducible gene *IFI44*, whose expression is regulated by IFN-γ exposure(38). Interestingly, *IFI44* has been suggested as a key target for controlling the cytokine storm-induced immunopathology observed in patents with influenza vi-rus and high pathogenic coronavirus infections(39). Based on our microarray expression analyses, we hypothesized that *ACE2* transcriptional regulation is associated with an immune-signaling pathway involving IRF and STAT factors.

An important limitation of microarray data concerning *ACE2* is the inability to discriminate between full-length *ACE2* and the recently-discovered short-length isoform *dACE2*(40). Therefore, the relative contribution of full-length ACE2 versus short-form dACE2 to these expression profiles remains unclear. We therefore sought more explicit, experimental interrogation of ACE2 gene regulation by considering the *cis-*regulatory landscape of the *ACE2* locus.

### Identification of functional intronic *ACE2* regulatory elements with *STAT1* and IRF1 binding sites

Gene expression is controlled by regulatory sequences bound by transcription factors. We next examined the regulatory region surrounding *ACE2*, compiling chromatin-accessibility datasets (i.e., Dnase-I Hypersensitivity Sites, (DHS)) from *in-vitro* and adult *in-vivo* lung samples from the ENCODE project (Figure 2). We identified six putative regulatory regions overlapping either cell-line or primary tissue DHS signals, a number of which also possess potential binding motifs for STAT1 and IRF1. We then refined this list to three putative regulatory elements within intronic regions of *ACE2* (Regions 1,4 and 5 in Figure 2) that overlap DHS data from both lung cell and tissue data and contained either predicted STAT1 and IRF1 binding motifs and/or aggregated ChIP-seq datasets for each factor (see Methods). These predicted factor binding motifs may be directly bound by STAT1 (Regions 1 and 5), and one possibly bound by IRF1 (Region 5).

**Figure 2.**
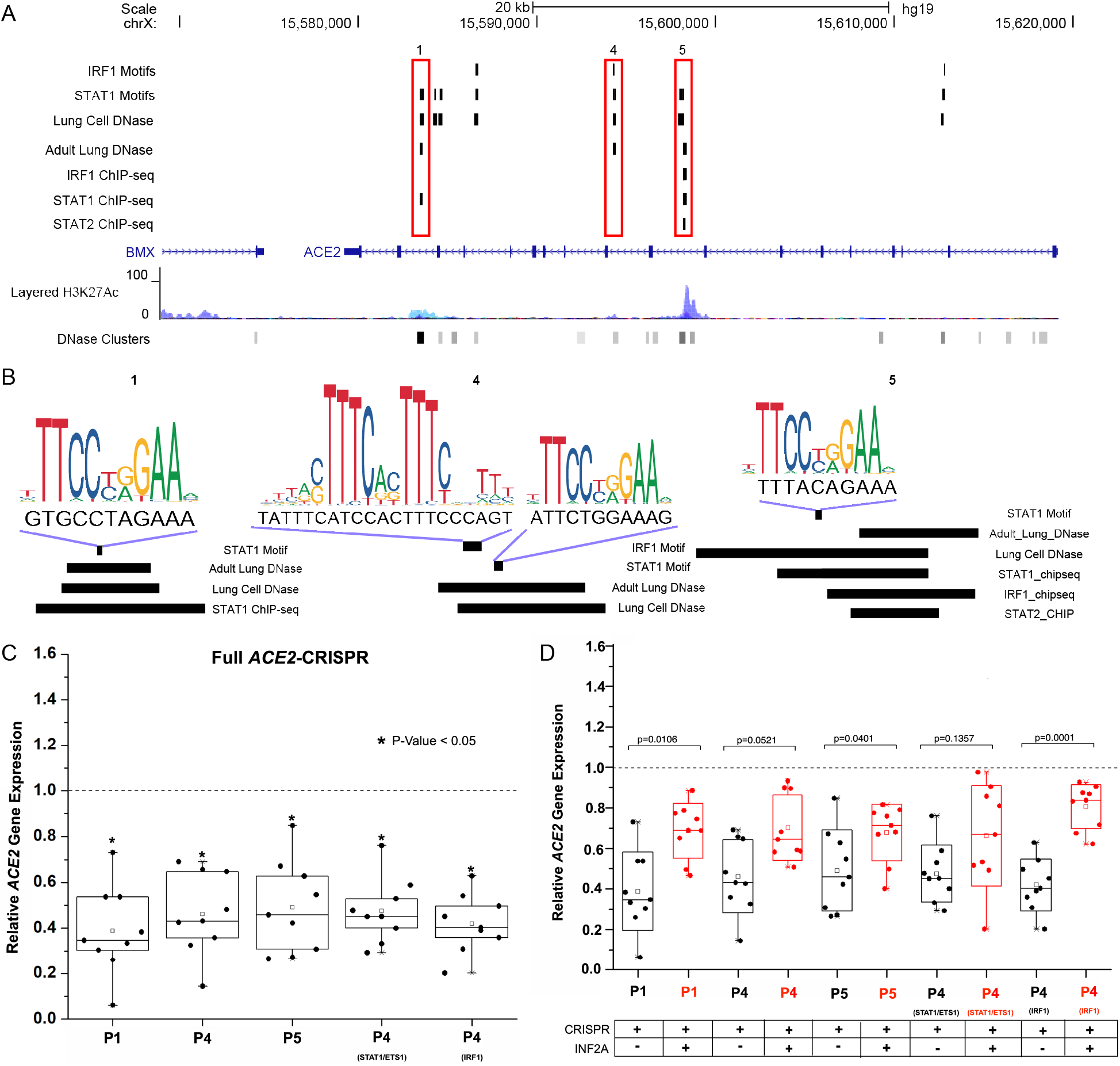
Identification of putative viral-response elements (STAT1 and IRF1 binding sites) in the *ACE2* intronic region. **a)** Identified transcription factor binding sites in *ACE2* intronic regions in the human genome (hg19). Three separate regions labeled 1,4,5 contain overlapping ChIP-seq peaks including IRF1, STAT1, and STAT2 binding sites, as well as DHS in lung cell lines indicative of open-chromatin and active transcriptional regulation. **b)** DNA sequence matches to predicted IRF and STAT transcription factor binding sites in the three regions identified above, with corresponding ChIP-seq peaks indicated as horizontal bars. **c)** Deletion of regions leads to decreased expression of full-length *ACE2*. **d)** Reductions in expression become attenuated when elements are deleted in the presence/absence of IFN-α treatment. See also Table S2.

In order to test the functionality of these three putative regulatory elements on *ACE2* regulation, we designed CRISPR guide RNAs (sgRNAs) to target and delete each element. We also designed sgRNAs to target predicted STAT1 and IRF1 motifs in Region 4 (Table S2). We tested our targeting strategy *in-vitro* on a human lung epithelial cell line (Calu-3). To rule out potential off-target effects, we first confirmed that transfection of sgRNA plasmids did not disrupt the expression of nearby genes. Specifically, expression levels of nearby *TMEM-27* and *BMX1* were not significantly altered with deletion of any element or putative binding site (Table S2). Using full-length isoform-specific primers, we next assessed levels of full-length *ACE2* transcripts using qPCR in wildtype and CRISPR-deleted cells. We found that full-length *ACE2* expression was significantly decreased with deletions of each individual element, as well as the targeted binding sites within Region 4 (Figure 2C, Table S2). A recent study identified that the dACE2 isoform is regulated upon SARS-CoV-2 infection(40). Interestingly, using primers specific to *dACE2*, we found that its transcript levels were also significantly decreased in our deletion experiments, and to a greater degree compared to full-length *ACE2* (Table S2). These results indicate that, in the absence of additional perturbation (i.e., above transfection), each of the three candidate intronic regulatory sequences we tested acts as an enhancer specifically for *ACE2*.

Our bioinformatic analyses of RNA-sequencing datasets and subsequent motif/ChIP-seq scans suggest that *ACE2* expression is regulated by an immune signaling pathway, possibly through STAT1 and IRF1 binding activity intronic to the *ACE2* locus. We therefore tested the effects of deleting these putative immune-responsive elements and specific binding sites in the context of immune signaling. Type I Interferons (IFNs), such as IFN-α are our first line of defence against invading viruses. We used IFN-α treatment to induce intra-cellular immune signaling pathways that would occur during viral infections(41) (see Methods). We first performed this experiment on wildtype cells and found that this treatment did not lead to significant increase of full-length *ACE2* transcripts (Figure S2a), although *dACE2* levels increased significantly, consistent with a previous study(40) (Table S2). We independently confirmed this finding at the protein level (Figure S2b), and further found that additional potent inducers of immune signalling, such as poly(I:C) treatment and direct infection with SARS-CoV-2, did not lead to significant up-regulation of full-length ACE2 at the transcriptional or protein levels.

We next performed IFN-α treatment in the context of enhancer deletion. We observed a significantly decreased effect of CRISPR element deletion on full length *ACE2* gene expression reduction with IFN-α stimulation compared to expression changes in the absence of stimulation (Figure 2d, Table S2). This was observed across the majority of our element and subelement (i.e., motif) deletions. This attenuated down-regulation was also observed for *dACE2* across stimulation-deletion experiments (Table S2). These findings suggest that these enhancer elements may be in part responsive to immunological stimulation (via IFNs) and play a role in a more complicated regulatory mechanism for *ACE2* expression (see Discussion).

### *ACE2* gene expression in lung epithelial cells is correlated with smoking and COPD disease status and associated with an *NRF2* antioxidant response

While much of the initial medical literature associated with SARS-CoV-2 patient demographics suggest a link between smoking status and disease severity(42–44), more recent studies have cast doubts as to the strength and significance of this relationship(45,46). The relationship between smoking history and respiratory viral infection disease severity has been suggested to be more complicated(47). It is also worth noting that ACE2, in addition to being the primary receptor for SARS-CoV-2 infection(5,7), serves an important biological role in multiple tissues(2), and is present in lung epithelium(3,4). Thus, shifts in basal expression levels of this protein, especially over time, may contribute to lung dysfunction in an indirect, more complex manner than can be measured using metrics such as COVID-19 disease severity. Given this possibility, we next assessed expression patterns of bronchial brushing datasets from current and previous smokers, focusing on *ACE2* and other co-expressed genes (e.g. *DUOX2*).

We analyzed a dataset of 159 healthy current smokers versus healthy former smokers, and identified the top 200 *ACE2* correlated genes (Figure 3, Table S1). Expression patterns for *ACE2* suggest current smoking status is associated with increased ACE2 levels, consistent with previous observations(48–50), and that this also accounts for the increased *ACE2* in COPD patients (Figure 3a). Expression patterns for *ACE2*-correlated genes alone were able to effectively distinguish smokers from non-smokers (Fig 3b). Functional enrichment analysis showed that in this dataset, genes co-expressed with *ACE2* are significantly associated with the NRF2 pathway, oxidative stress, glutathione metabolism, and TGF-β regulation of the extracellular matrix. *NRF2* is a key transcription factor that regulates the oxidative stress response in the lung(51,52). Consistent with this, according to both ChIP-seq data and gene expression perturbation data, *NRF2* (*NFE2L2*) was the top transcription factor identified as a likely regulator of these genes;; for example, the NRF2-regulated antioxidant gene *NQO1* was the fourth ranked *ACE2* correlated gene in this dataset. *ACE2* correlated genes also overlapped significantly with genes upregulated by the transcription factor ETS1 (GSE21129)(53);; ETS1 is an important regulator of ROS in response to angiotensin II, linking it to ACE2 function(54). Moreover, *ETS1* expression is induced by reactive oxygen species (ROS) exposure through an antioxidant response(55). Thus, *ACE2* expression in smokers appears to be associated with oxidative stress gene regulation, likely mediated by NRF2 and ETS1.

**Figure 3.**
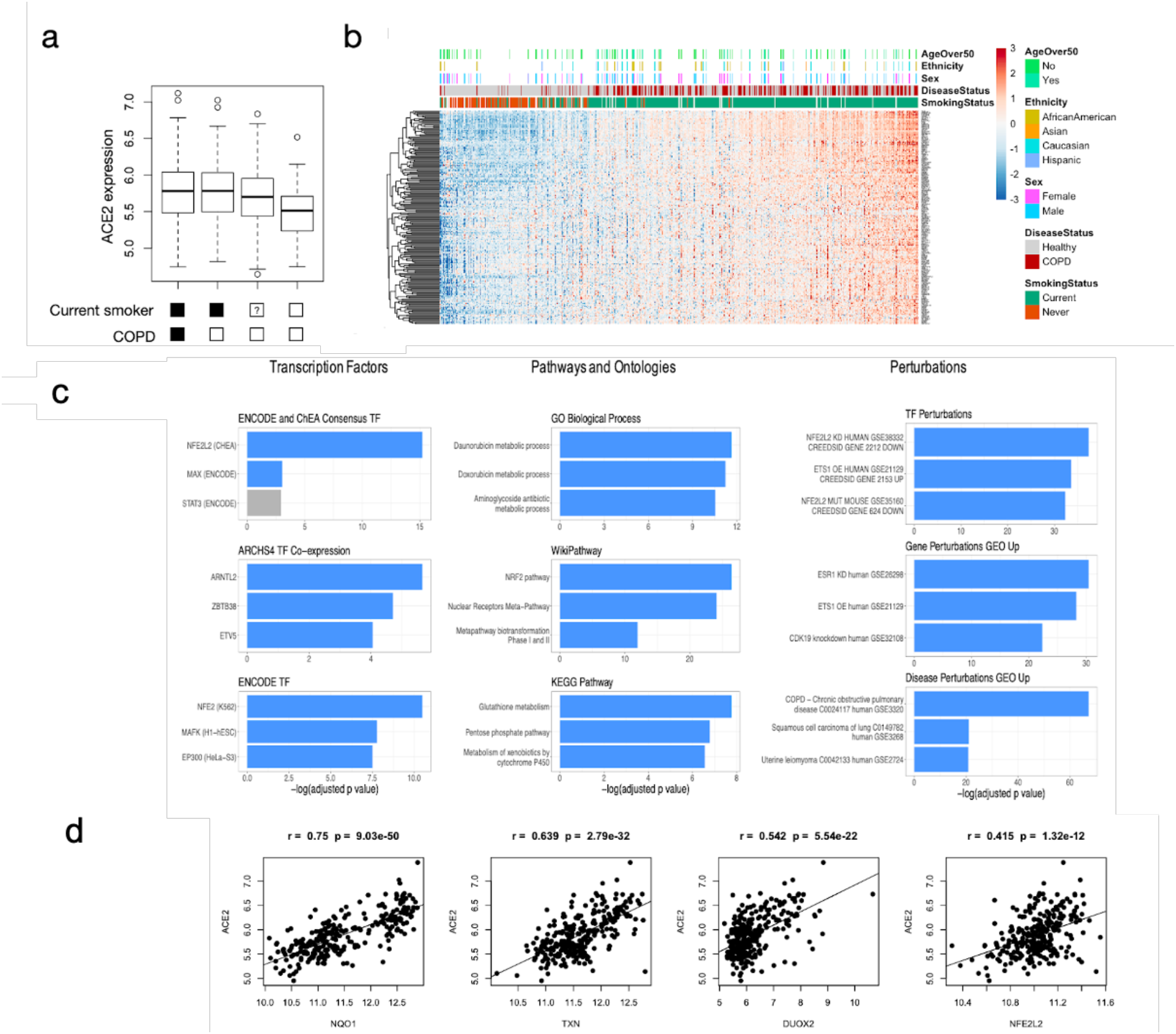
Expression and functional enrichment analysis of *ACE2* and co-expressed genes in smokers and individuals with COPD. **a)** Analysis of relative *ACE2* expression with respect to smoking status and COPD diagnosis. **b)** Expression of top 200 *ACE2*-correlated genes (including *ACE2*) in individuals with various smoking status and COPD diagnosis (N=159). **c)** Functional enrichment analysis of top 200 *ACE2*-correlated genes (including *ACE2*). Terms are ranked by -log_2_(FDR-adjusted *p* value) for nine ontologies/groups of interest. **d)** Pearson correlation of *ACE2* with important interferon-related candidate genes found to be co-expressed with *ACE2*.

To verify these trends, we repeated the same analyses with a second cohort dataset from a different microarray platform associated with 345 healthy smokers versus healthy non-smokers (Figure S3). Notably, this dataset consists predominantly of younger individuals (age < 50), whereas the first dataset includes predominantly older individuals (> 50). *ACE2* co-expression patterns were highly correlated between the two independent datasets providing support that these are robust signals. As with the first analysis, genes co-expressed with *ACE2* showed the strongest associations with *NRF2* gene targets. Moreover, *NRF2* and *ETS1* formed the top three overlapping datasets according to enrichments for TF Perturbation datasets (Figure S3c). This dataset also included a larger number of COPD patients;; that we observed similar patterns of *ACE2* expression and co-expressing genes in this analysis may suggest similar effects on smoking and disease status on *ACE2* regulation (see Discussion).

### Identification of functional intronic *ACE2* regulatory elements with possible antioxidant-response element (ARE) activity

Our microarray data analyses led to a predicted association between oxidative stress and *ACE2* levels, which prompted us to consider the existence of antioxidant-response elements (AREs) within the *ACE2* locus (Figure 4a). We performed an unbiased analysis of the locus and regulatory regions intronic to *ACE2* (Figure 4b). Our analysis identified AP-1 as the top enriched transcription factor binding site in the locus, suggesting that AP-1 may be an additional regulator of *ACE2*. Klatt *et al*. have shown AP-1-c-Jun subunit binding to DNA is dependent on the cellular GSH/GSSG ratio, a marker of cellular ROS levels(56). Looking at the top *ACE2* correlated genes across all datasets, we identified significant co-expression between *ACE2* and *FOSL2*, as well as *ACE2* and *JUN*. This suggests that AP-1 may be a transcription factor involved in oxidative stress-mediated regulation of *ACE2* levels.

**Figure 4.**
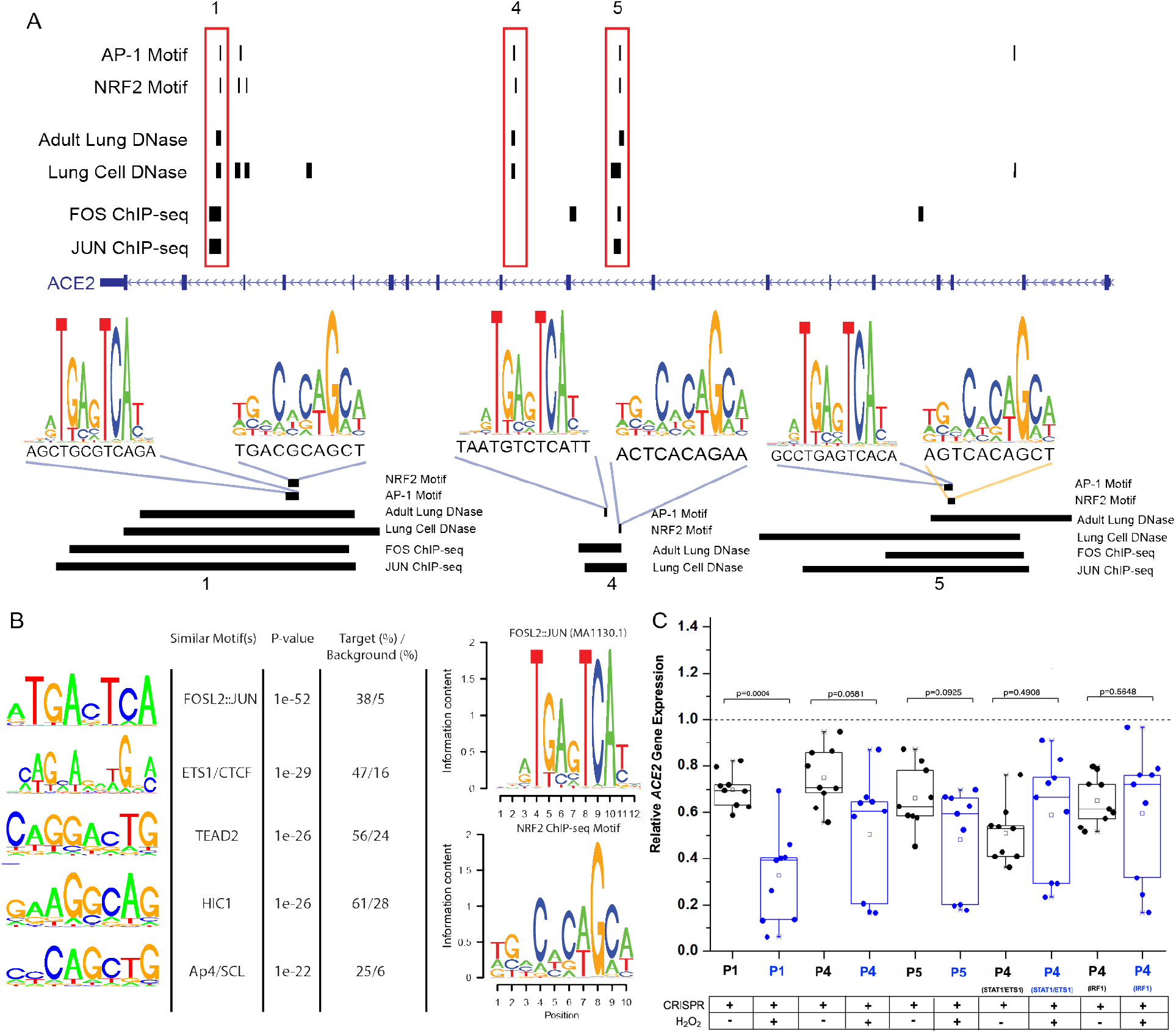
Putative anti-oxidant response elements (AREs) in the *ACE2* regulatory region. **a)** Predicted AP-1 and NRF2 binding sites in *ACE2* intronic regions in the human genome. Three separate regions labeled 1, 4, and 5 (red boxes) contain overlapping ChIP-seq peaks including FOS/JUN binding sites, as well as DNase hypersensitivity peaks in adult lung tissue and cell line samples indicative of open-chromatin and active transcriptional regulation. Shown below are DNA sequence matches of predicted AP-1 and NRF2 transcription factor binding sites in the three regions identified above, with corresponding ChIP-seq peaks indicated as horizontal bars. **b)** (Left) Top statistically over-represented motifs in *ACE2* non-coding regulatory regions and their top matches to known transcription factor binding preferences. The top two binding motifs identified bear strong resemblance to FOSL2:JUN (AP1) and (ETS1/CTCF). (Right) The JASPAR-database FOSL2::JUN binding motif (MA1130.1) was enriched in intragenic *ACE2* elements. Also shown is an NRF2 binding motif defined using ChIP-seq data. Both motifs in (**b**) were used to scan intragenic *ACE2* element sequences in (**a**). **c)** Deletion of regions leads to decreased expression of full-length *ACE2* in the presence/absence of oxidative stress (blue boxes). See also Table S2.

Given the observed motif enrichments for AP-1 and NRF2 transcription factor binding sites in the *ACE2* locus, we next looked at the individual predicted motif hits for these factors within the six putative regulatory regions defined above (Figure 4b). Three of these enhancers (Regions 1, 4, and 5) contained predicted AP-1 and NRF2 binding sites and were active in both *in-vivo* and *in-vitro* lung datasets. Two of these regions (Region 1 and 4) also overlapped ChIP-seq datasets for FOS and JUN factors, important co-factors associated with AP-1 complex(57) and NRF2(58) enhancer binding, respectively. We next sought to delete these elements in the context of oxidative stress, with the expectation that, if these elements act as AREs, that the effects of deletion should be magnified during an ROS response.

We first examined *ACE2* transcript levels after exposing wildtype Calu-3 cells to hydrogen peroxide (0.5mM), a potent ROS(34) (see Methods). We found that exposure to hydrogen peroxide led to significant decreases in expression of both full-length *ACE2* and *dACE2* (Figure S4, Table S2). Previous mouse studies have also demonstrated a down-regulation of *ACE2* levels and activity following acute ROS exposure(59). However, *ACE2* levels are up-regulated with chronic oxidative stresses(51,60);; this may suggest a more complicated regulation of *ACE2*, possibly as a function of time (see Discussion).

We next deleted each regulatory element and assessed *ACE2* expression in the presence/absence of hydrogen peroxide treatment (see Methods). We observed a significant decrease in basal levels of full-length *ACE2* transcripts for the majority of regions/sites deleted in the absence of external stimulus (Table S2). In the context of exogenous oxidative stress, we observed a further significant decrease only for Region 1, whilst all other trended downwards (Table S2). For this first region, the magnitude of down-regulation was significantly greater under oxidative stress when compared to the unstimulated deletion change (Figure 4c), while for other regions magnitudes were similarly larger despite the lack of significance (potentially due to the increased variability in expression observed in ROS-stressed cells). This first region contains predicted NRF2 and AP-1 motifs, and also overlaps with both FOS and JUN ChIP-seq signals, possibly explaining the increased effect of deletion under ROS conditions. Finally, we again saw these differences to be accentuated when considering levels of the *dACE2* transcript (Table S2).

## Discussion

Regulation of *ACE2* expression at the transcriptional level may impact susceptibility to viral infection. Subsequently, changes to *ACE2* expression during viral infection can lead to an imbalance in renin-angiotensin system (RAS) signalling contributing to the manifestation of clinical symptoms such as excessive inflammation(61), myocardial injury(10), and lung injury(62). Indeed, it has recently been suggested that targeting transcriptional inhibition of *ACE2* expression may be a therapeutic avenue for prevention of severe COVID-19 infection(63), while counteracting infection-induced *ACE2* down-regulation is being pursued as a therapeutic treatment to reduce disease severity(11).

In the first part of our study, we utilized microarray expression datasets from healthy, non-smokers and identified genes whose expression patterns significantly correlated with that of *ACE2*. Groups of correlated genes may suggest shared upstream regulators;; gene-set enrichment analyses indicated that *ACE2* and correlated genes may be under the control of immune signalling pathways integrating on the STAT and IRF families of transcription factors – namely, STAT1 and IRF1. These results were also observed when performing a separate analysis on asthmatics individuals (Figure S1). These factors have also previously been associated with possible epigenetic regulation of *ACE2* expression.

Given these findings, we considered *cis*-regulatory elements in the *ACE2* locus which may be proximate mediators of this immune-response regulatory mechanism. After identifying six such regions, we prioritized and then experimentally deleted three putative intronic enhancers, containing predicted STAT1 and IRF1 binding motifs. Deletion of these three elements (Regions 1, 4, and 5) lead to a consistent down-regulation of *ACE2* transcripts. The down-regulation of *ACE2* upon deletion, relative to mock-transfected controls, suggests that these enhancers may contribute to basal expression. Interestingly, performing these deletions in the context of immune stimulation caused a significant attenuation of this effect, while immune stimulation in wildtype cells caused no significant changes to expression (Figure S2). These latter results corroborate our previous findings that SARS-CoV-2 infection does not up-regulate transcript levels of *ACE2* in spite of significant increases in type I and type III IFNs, as well as up-regulation of known interferon-stimulated genes (e.g. *IFIT1, IRF7, OAS2*)(31). The observed attenuation effect may suggest an upper-threshold or ‘saturation’ of *ACE2* expression from baseline, such that immune signalling does not lead to a significant increase. However, following deletion of these putative enhancers, proximate regulators of immune signaling (e.g. STAT1, IRF1) acting elsewhere in the *ACE2* locus (e.g., at the promoter level) may be able to compensate for the loss/reduction in enhancer activity. Further experimentation (e.g., using a viral-infection model system) may elucidate the role that these enhancers play in up-regulating *ACE2* during an immune response.

An understanding of the mechanisms regulating *ACE2* expression during viral infection is important from a disease-pathology point of view, given that this may inhibit the protective effects of ACE2 activity. In addition, understanding the regulatory mechanisms controlling *ACE2* expression prior to viral exposure may be of equal importance from a disease-prevention point of view, given that baseline levels of *ACE2* in high-exposure tissues (e.g., lung) may modify viral susceptibility(14). It has been suggested, though not conclusively shown, that chronic smokers are at elevated risk to both SARS-CoV-2 infection as well as severe disease(42,44,46). This follows with previous studies of other coronaviruses, e.g. MERS-CoV, for which epidemiological evidence does suggest smoking status as a key risk factor(64,65). In terms of increased susceptibility, it may be that smokers have elevated base-line *ACE2* expression in lung tissues, increasing the likelihood that SARS-CoV-2 may bind their target receptors(63).

In the second part of our study, we analyzed microarray expression data from two independent datasets consisting of current smokers, non-smokers, and COPD patients. Considering the expression of *ACE2*, we also observed previously-reported increases in baseline expression within smokers(60,63). With this, as well as genes showing similar transcriptional behaviours, we identified enrichments for oxidative stress-response pathways, including transcriptional regulators such as NRF2 and the AP-1 complex. These signals are indicative of another potential regulatory mechanism acting on the *ACE2* locus, and are expected given the chronic oxidative stresses experienced by habitual smokers and patients with COPD(66).

Looking again at the *ACE2* locus we found enrichment in open-chromatin regions (putative enhancers) for DNA sequences bearing similarity to known FOSL2::JUN binding motifs, further suggesting the regulatory effects of oxidative-response signalling at this locus. We therefore performed another set of targeted deletion experiments of putative intronic enhancers most likely to behave as antioxidant-response elements (AREs) (e.g. contain NRF2 motifs, AP-1 ChIP-seq signals, etc.). Performing these deletions in the context of exogenous oxidative stress yielded a substantial decrease in *ACE2* expression for the first element tested, with a fold-change decrease below that observed in wildtype cells following treatment.

We suggest that this putative ARE, and potentially others which trend in the same direction, act to counter the inhibitory effects of oxidative stress on *ACE2* expression, which has been previously observed in a mouse model of hyperoxia(59), preventing a more deleterious loss of ACE2 protein following acute exposure. ACE2 plays an important role in mitigating acute oxidative stress(67,68), particularly in the context of cardiovascular and lung disease(69,70). We further propose that the repression of *ACE2* upon acute oxidative stress, when repeated on the order of decades in chronic smokers, may lead to an ‘over-compensation’ of baseline *ACE2* expression – establishing higher levels of ACE2 protein to protect lung tissues from further damage. This process could be mediated by a number of oxidative stress-response mechanisms;; in particular, our observed enrichments for NRF2-regulated genes co-expressed with *ACE2*, along with the presence of NRF2 motifs within intronic *ACE2* enhancers, follows with the protective role of NRF2 signaling induced in response to cigarette smoke(71). Furthermore, a mouse-model study of cigarette smoke found significant increases in ACE2 activity only after three weeks of exposure(72), while additional studies have found dose-response effects with increased treatment time(51,73). Human smokers also exhibit a dose-response effect of *ACE2* expression with increasing pack-years(60). However, we acknowledge the speculative nature of this proposed over-compensating effect and note the importance of additional experimental testing. While the links between smoking status and COVID-19 severity are controversial, the link between COPD status and COVID-19 severity may be clearer(47,74,75). More generally, it has been suggested that the detrimental effects of smoking, most notably attenuation of antiviral innate immune responses(76), can increase susceptibility to pathogen infection(65).

In this study, we explored the regulatory mechanisms which may act on the *ACE2* locus in the context of both immune stimulation as well as oxidative stress, leading us to identify two putative pathways which may mediate this transcriptional regulation. It is important to note that these pathways are not mutually-exclusive;; the links between immune signalling and oxidative stress are well-established(77), and this is particularly true for ACE2 given its biological role in RAS regulation(2,5). We suggest that further experimental testing is warranted to confirm these predicted mechanisms, and furthermore to develop potential strategies taking advantage of this knowledge to modify susceptibility and disease severity of coronavirus infections, particularly SARS-CoV-2.

## Materials and Methods

### *ACE2* co-expression and functional enrichment analysis with public microarray data

Public microarray experiments using Affymetrix chips (HuGene-1.0-st-v1 and HG-U133 Plus 2) on airway epithelial cell samples were obtained from the NCBI Gene Expression Omnibus (GEO) database, as described in our previous work(4). This resulted in a total of 1859 individual samples from 33 different experiments (See Table S1). Within this dataset, disease status (Healthy: 504, COPD: 338, Asthma: 136) and/or smoking status (Never: 409, Former: 139, Current: 956) information was included for 1716 samples. For all datasets, raw intensity values and annotation data were downloaded using the *GEOquery* R package (version 2.52.0)(16) from the Bioconductor project. Probe definition files were downloaded from Bioconductor and probes were annotated using Bioconductor’s *annotate* package. All gene expression data were unified into a single dataset that was then RMA-normalized, and only genes present in both of the Affymetrix platforms (N = 16,105) were kept for subsequent analyses. Correction of experiment-specific batch effects was performed using the ComBat method implemented using the *sva* R package (version 3.32.1).

Top *ACE2* co-expressed genes were identified based on the 200 highest Pearson correlation (*r*) values to *ACE2*. Heat maps for top 200 *ACE2*-correlated genes across samples were generated with the *pheatmap* R package (version 1.0.12). For display only, expression values were row-normalized (across gene) using the ‘scale’ function in base R, and converted to Z-scores. For heat map coloring, a “ceiling” and “floor” of +3 and -3 was applied to the Z-scores. Histograms and *ACE2* correlation scatter plots were generated with the *base* R package (version 3.6.3)(17). Gene lists are available in Table S1.

The top 200 *ACE2*-correlated genes were analyzed using Enrichr(18) to identify enriched pathways and functional ontologies. Terms were ranked within ontologies by FDR-adjusted *p* value (calculated by Enrichr by running the Fisher exact test for random gene sets in order to compute a mean rank and standard deviation from the expected rank for each term in the gene set library) and the top 3 terms for ontologies of interest were selected. Functional enrichment bar plots were generated with the *ggplot2* R pack-age (version 3.2.1).

### *ACE2* regulatory region analyses

ENCODE(19) DNase-seq datasets were obtained for adult lung (File accessions: ENCFF271JAF, ENCFF331SYD, ENCFF681UOZ) and primary cell (ENCFF165ZIA, ENCFF334RSR, ENCFF338GII, ENCFF446FTN, ENCFF460ZFL, ENCFF546QUZ, ENCFF644XOI, ENCFF957JQC) samples as processed hg19 bed-formatted files. Called peaks (putative regulatory elements) were subsequently intersected within lung/cell sets using *bedtools*(20) (version 2.26.0), requiring that a peak be replicated in at least two samples for retention. To capture a broader regulatory region around the *ACE2* gene, peaks falling within 1MB upstream/downstream of the *ACE2* promoter were collected and pooled across lung/cell sets. Human hg19 reference sequences were obtained for these elements using UCSC(21).

Sequences were subsequently used for *de novo* motif analysis using HOMER (version 4.10)(22) using a 10x random shuffling as a background set. *De novo* motifs were compared to a vertebrate motif library included with HOMER which incorporates the JAS-PAR(23) database. Matches are scored using Pearson’s correlation coefficient of vectorized motif matrices (PWMs), with neutral frequencies (0.25) substituted for non-overlapping (e.g., gapped) positions. Best-matching motif PWMs obtained from this analysis are shown in Figure 4B. The highest-rank motif bore close similarity to the *FOSL2::JUN* PWM from the JASPAR database (MA1130.1). This reference PWM was subsequently used for motif scanning. The AME program(24), part of the MEME-Suite(25), was also used with these sequences to look for enrichments of known transcription factor (TF) motifs. Focusing on elements intragenic to *ACE2* as more proximate candidate regulatory regions, these reference sequences were also tested for enriched motifs using AME. The *FOSL2::JUN* (MA1130.1) PWM was used to scan these sequences using TFBSTools(26) (version 1.24.0). Hits of minimum sequence scores of 80% were retained, and subsequently filtered for Benjamini-Hochberg adjusted *p* value < 0.05. For illustrative purposes, the best-scoring hit for each hit-containing element was selected. JASPAR-database motif matrices were also obtained for *STAT1* (MA0137.3), and *IRF1* (MA0050.2) and similarly used to scan intragenic *ACE2* elements using TFBSTools as described. Given the observed expression data for *NRF2*, a ChIP-seq dataset for this factor (GSE37589)(27) was downloaded as called hg18 peaks. These were lifted-over to hg19 using the UCSC liftOver utility(21) with sulforaphane and vehicle-treatment datasets pooled and merged for a final set of 919 peaks. Reference hg19 sequences were obtained for these peaks and used with HOMER to define an *NRF2 de novo* motif;; the resulting PWM was subsequently used with TFBSTools to scan the putative *ACE2* intragenic regulatory sequences as described above. ChIP-seq data for indicated factors (*IRF1, STAT1, STAT2, FOS* and *JUN*) were obtained from ChIP-ATLAS(28) as an aggregate across all cell types using a significance threshold of 50 (*q* value < 1e-5). Peak files (hg19) were sorted and merged for overlapping peaks using bedtools. DNase, ChIP-seq, and motif hit tracks were loaded into the UCSC Genome Browser(29) for visualization.

### *In-vitro* experimental validation studies

#### Poly(I:C) transfection and IFN treatment

Calu-3 cells were mock transfected with 4 μL of lipofectamine 3000 (ThermoFisher Scientific) in Opti-MEM (ThermoFisher Scientific) only or transfected with 100 ng of poly(I:C) (InvivoGen) for 6 hours. Recombinant human IFNβ1 was generated using *Drosophila* Schneider 2 (S2) cells following manufacturer’s recommendation and by using ThermoFisher Scientific’s *Drosophila* Expression system (ThermoFisher Scientific). Recombinant IFNβ1 was collected as part of the cell culture supernatant from S2 cells and total protein was measured using Bradford assay. Total protein concentration was used for subsequent experiments. To demonstrate that S2 cell culture media did not contain non-specific stimulators of mammalian antiviral responses, we also generated recombinant green fluorescent protein (GFP) using the same protocol and used the highest total protein concentration (2 mg/ml) for mock treated cells. SARS-CoV-2 infected cells were treated with supernatant containing IFNβ1 or GFP for 6 hours.

#### SARS-CoV-2 infection

Clinical isolate of SARS-CoV-2 (SARS-CoV-2/SB3) was propagated on Vero E6 cells and validated by next generation sequencing(30). Virus stocks were thawed once and used for an experiment. A fresh vial was used for each experiment to avoid repeated freeze-thaws.

#### Immunoblots

Calu-3 cells were seeded at a density of 3 × 10^5^ cells/well in 12-well plates. Cells were infected with SARS-CoV-2 at an MOI of 1. Control cells were sham infected. Twelve hours post incubation, cells were transfected or treated with poly(I:C) or IFNβ, respectively for 6 hours. Cell lysates were harvested for immunoblots and analyzed on reducing gels as mentioned previously(31,32). Briefly, samples were denatured in a reducing sample buffer and analyzed on a reducing gel. Proteins were blotted from the gel onto polyvinylidene difluoride (PVDF) membranes (Immobilon, EMD Millipore) and detected using primary and secondary antibodies. Primary antibodies used were: 1:1000 mouse anti-SARS/SARS-CoV-2 N (ThermoFisher Scientific;; Catalogue number: MA5-29981;; RRID: AB_2785780), 1:1000 rabbit anti-beta-actin (Abcam;; Catalogue number: ab8227;; RRID: AB_2305186), and 2 μg/ml of mouse anti-ACE2 (R&D Systems;; Catalogue: MAB933;; RRID: AB_2223153). Secondary antibodies used were: 1:5000 donkey antirabbit 800 (LI-COR Biosciences;; Catalogue number: 926-32213;; RRID: 621848) and 1:5000 goat anti-mouse 680 (LI-COR Biosciences;; Catalogue number: 925-68070;; RRID: AB_2651128). Blots were observed and imaged using Image Studio (LI-COR Biosciences) on the Odyssey CLx imaging system (LI-COR Biosciences).

#### Cell line and culture condition

The human lung adenocarcinoma cell line, Calu-3 (ATCC HTB-55) were grown in Minimum Essential Medium (MEM)-α (Gibco, Gaithersburg, Maryland) supplemented with 10% fetal bovine serum (FBS), and 1% penicillin-streptomycin (P/S) in 5% CO_2_ at 37°C. Media was replaced every 2 days and the cells were subcultured every 5 days. The cells were passaged and used in experimental assays without additional STR authentication or mycoplasma testing.

#### CRISPR targeting of *ACE2* regulatory elements *in vitro*

All sgRNAs flanking human *ACE2* regulatory elements and sub-elements containing TF binding sites were designed using the Genetic Perturbation Platform (GPP) sgRNA design tool from Broad Institute (https://portals.broadinstitute.org/gpp/public/), synthesized by Integrated DNA Technologies, Inc (Coralville, Iowa), and cloned into the PX458 vector following published protocols(33). The sequence of all sgRNAs along with their chromosomal locations (hg19) are listed in Table S2.

Guide RNAs (see Table S2), flanking the *ACE2* regulatory elements and sub elements containing TF binding sites, were first tested for the ability to induce efficient deletions of the enhancer elements/sub elements in cultured Calu-3 cells (N = 3 biological replicates per assay). Calu-3 cells were maintained in MEM α media supplied with 10% FBS (Gibco) and 1% Pen/Strep (0.025%) and seeded in a six-well plate for 1-day prior to transfection. After culturing at 37°C with 5% CO_2_, the cells were scanned for GFP fluorescence under GFP-microscope at 24 h to verify successful transfection efficiency (i.e., >70% of the cells were GFP positive). After 48 h of CRISPR experiment, DNA was extracted from the CRISPR-cas9 targeted Calu-3 cells using E.Z.N.A Tissue DNA Kit (Omega Bio-Tek, Norcross, GA), and the *ACE2* regulatory element/sub-element region was amplified using PCR primers flanking each sgRNA location (listed in Table S2). PCR amplified products were purified from 1% agarose gel using E.Z.N.A Gel Extraction Kit (Omega Bio-Tek, Norcross, GA). Sanger sequencing was used to verify successful deletion of the target region. To examine effects on *ACE2* and nearby gene expression (*TMEM-27* and *BMX1*), RNA was extracted from control and CRISPR-Cas9 targeted Calu-3 cells (N = 3 biological replicates, with 3 technical replicates per experiment per condition) and prepared using Trizol Reagent (Thermo Fisher Scientific, Springfield Township, New Jersey) and Direct-zol(tm) RNA Miniprep kit (ZYMO). Two micrograms of total RNA were used to synthesize first-strand cDNA using SuperScript III First-Strand Synthesis System (Thermo Fisher Scientific). qRT-PCR analysis was then performed with gene specific primers and Applied Biosystems Power SYBR master mix (Thermo Fisher Scientific) with *ACTB* house-keeping gene as an internal control. sgRNAs and primers used for qRT-PCR are listed in Table S2.

#### CRISPR deletion of *ACE2* regulatory elements under interferon treatment

Briefly, the Calu-3 cells were cultured in MEM α media supplemented with 10% FBS (Gibco) and 1% Pen/Strep (0.025%), plated in 6-well plates and utilized at ∼70% confluence. The cells were then subjected to CRISPR-Cas9 mediated deletion of *ACE2* regulatory elements/sub-elements in the presence/absence of recombinant protein of human interferon, alpha 2 (IFNA2) (OriGene Technologies Inc, Atlanta, GA) (100 ng/ml) for 48 h. Following CRISPR-deletion under interferon treatment, DNA and RNA were extracted from the Calu-3 cells and used for genotyping and gene expression analysis, respectively.

#### CRISPR deletion of *ACE2* regulatory elements under H_2_O_2_ treatment

The Calu-3 cells were cultured in MEM α media as described above and subjected to CRISPR-cas9 mediated deletion of *ACE2* regulatory elements/sub elements for 48 h. CRISPR-cas9 targeted Calu-3 cells were treated with hydrogen peroxide (H_2_O_2_) using the protocol described previously(34). Following CRISPR deletion of *ACE2* regulatory elements/sub elements, the Calu-3 cells were challenged with or without oxidative stress by exposure to 0.5 mM (initial dose) of H_2_O_2_. As Calu-3 cells rapidly metabolize H_2_O_2_ in 1 h, H_2_O_2_ treatments were performed for 2 h, with additional bolus of H_2_O_2_ every 60 min for times longer than 1 h. DNA and RNA were extracted from the CRISPR-cas9 targeted Calu-3 cells subjected to H_2_O_2_ treatment, and used for genotyping and gene expression analysis, respectively.

## Acknowledgements

The authors would like to thank Harvard University Dean Christopher Stubbs and Dean Francis Doyle for granting restricted access to the Capellini Laboratory during the COVID-19 pandemic to perform these studies. The authors would like to thank members of the Capellini, Doxey, and Hirota labs for critical insight into this work. AB was the recipient of a fellowship from the Natural Sciences and Engineering Research Council of Canada (NSERC). VIDO receives operational funding for its CL3 facility (InterVac) from the Canada Foundation for Innovation through the Major Science Initiatives. VIDO also receives operational funding from the Government of Saskatchewan through Innovation Saskatchewan and the Ministry of Agriculture. ACD is funded in part by an NSERC Discovery Grant (RGPIN-2019-04266). TDC is funded in part by The American School of Prehistoric Research, Harvard University.

## Supplementary Tables

**Table S1: Microarray expression analyses**. Sheet 1: Metadata for microarray expression datasets downloaded and processed for this study. Sheet 2: The top 200 genes coexpressed with *ACE2* expression (by Pearson correlation) across the indicated expression analysis (relevant main/supplemental figure also indicated). Sheet 3: enrichR geneset enrichment results for gene sets shown in Sheet 2 – statistical results correspond with enrichments visualized in indicated main/supplemental figures.

**Table S2: *ACE2* regulation and experimental data**. Sheet 1: CRISPR Guide and PCR primer information. Sheet 2: Gene expression data for genes nearby *ACE2* following deletion of the indicated regulatory region. ‘VC’ – comparisons of deletion to vector-control samples (see Methods). Sheet 2: Expression of full-length *ACE2*, and the dACE2 isoform, following either IFN treatment of H_2_O_2_ treatment – results correspond to Figures S2 and S4, respectively. Sheet 3: Expression of full-length *ACE2*, and the dACE2 isoform, following deletion of the indicated region, either in the presence or absence of IFN treatment. Sheet 4: Expression of full-length *ACE2*, and the dACE2 isoform, following deletion of the indicated region, either in the presence or absence of H_2_O_2_ treatment.

## Supplementary Figures

**Figure S1.**
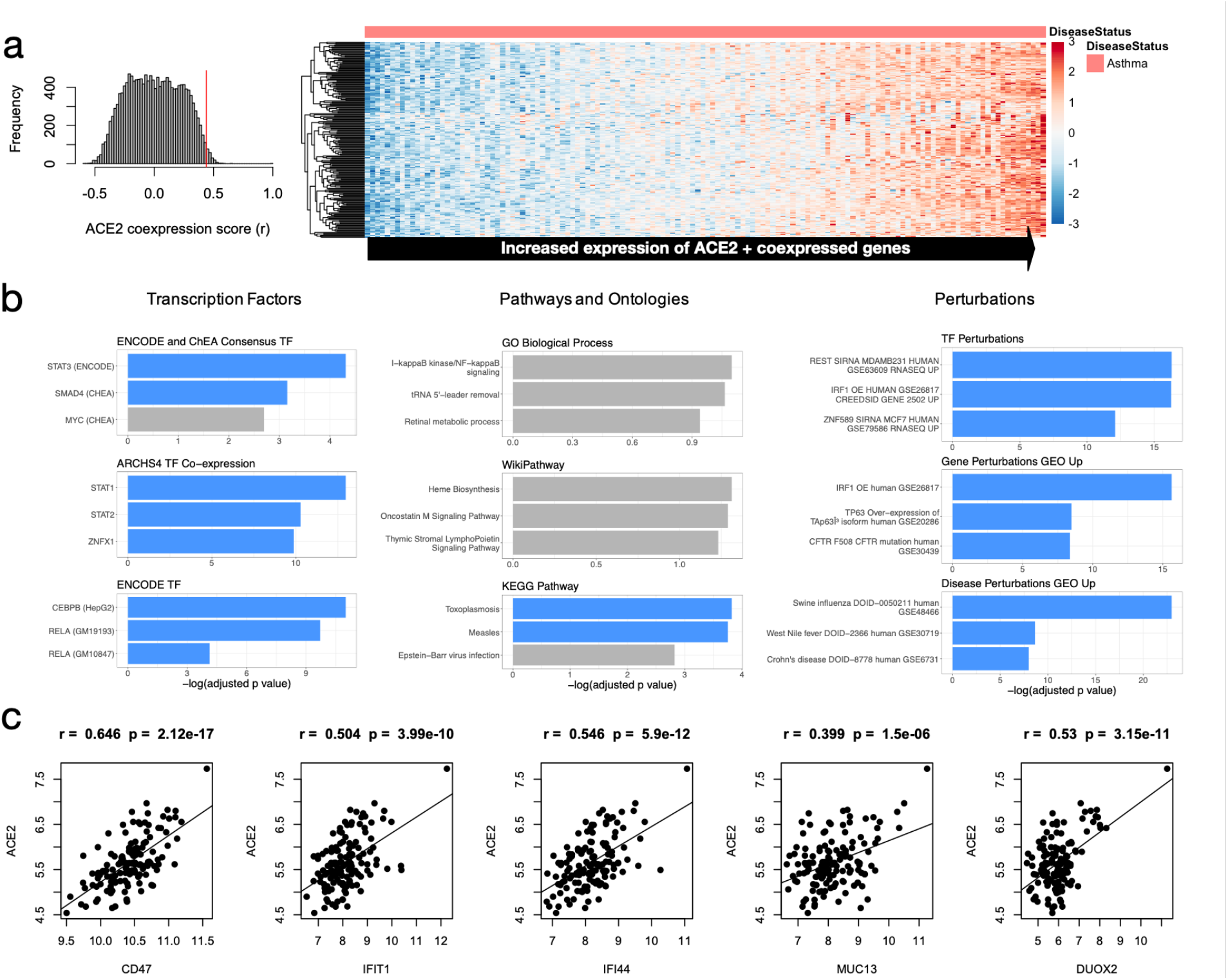
*ACE2* co-expressed genes and functional associations in asthmatics. **a)** Expression of top 200 *ACE2*-correlated genes (including *ACE2*) in asthmatics (N=136). **b)** Functional enrichment analysis of top 200 *ACE2*-correlated genes (including *ACE2*). Terms are ranked by -log_2_(FDR-adjusted *p* value) for nine ontologies/groups of interest. **c)** Pearson correlation of *ACE2* with important interferon-related candidate genes found to be co-expressed with *ACE2*.

**Figure S2.**
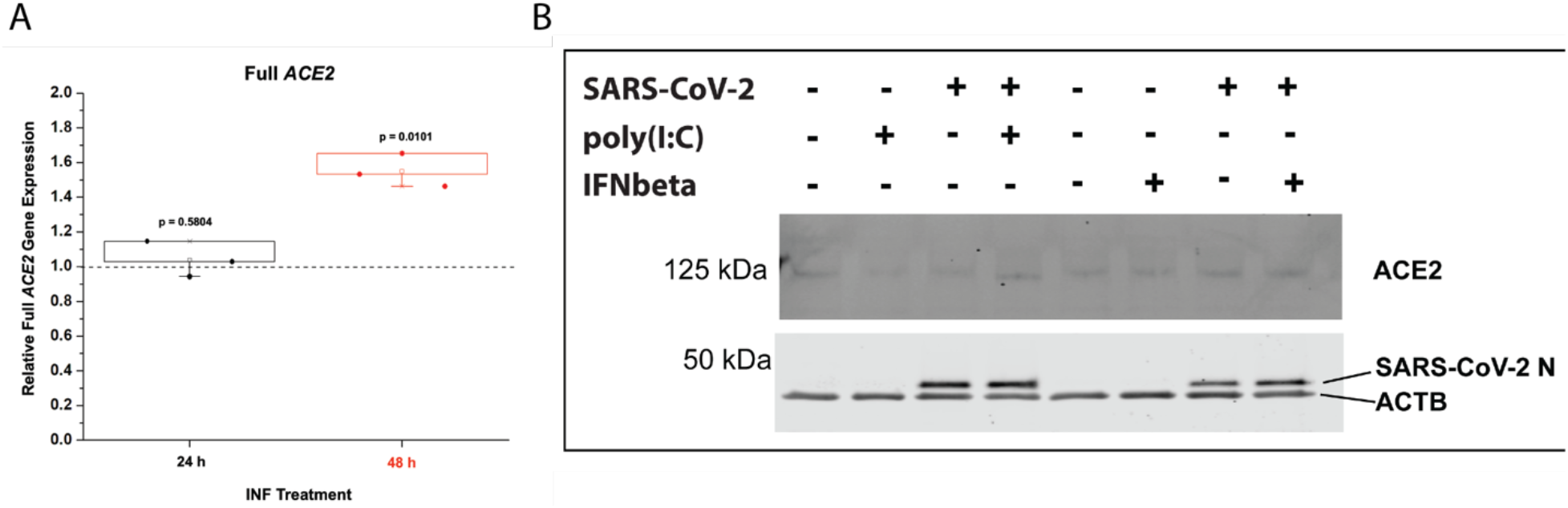
*ACE2* expression in response to immune signalling. (**A**) Expression of full-length *ACE2* following 24 or 48h of IFN-α treatment – p-values of expression change relative to untreated controls. (**B**) Immunoblots for ACE2 protein in Calu3 cell lysate in the context of SARS-CoV-2 infection, poly(I:C) treatment, and/or IFNβ1 treatment. See Methods.

**Figure S3.**
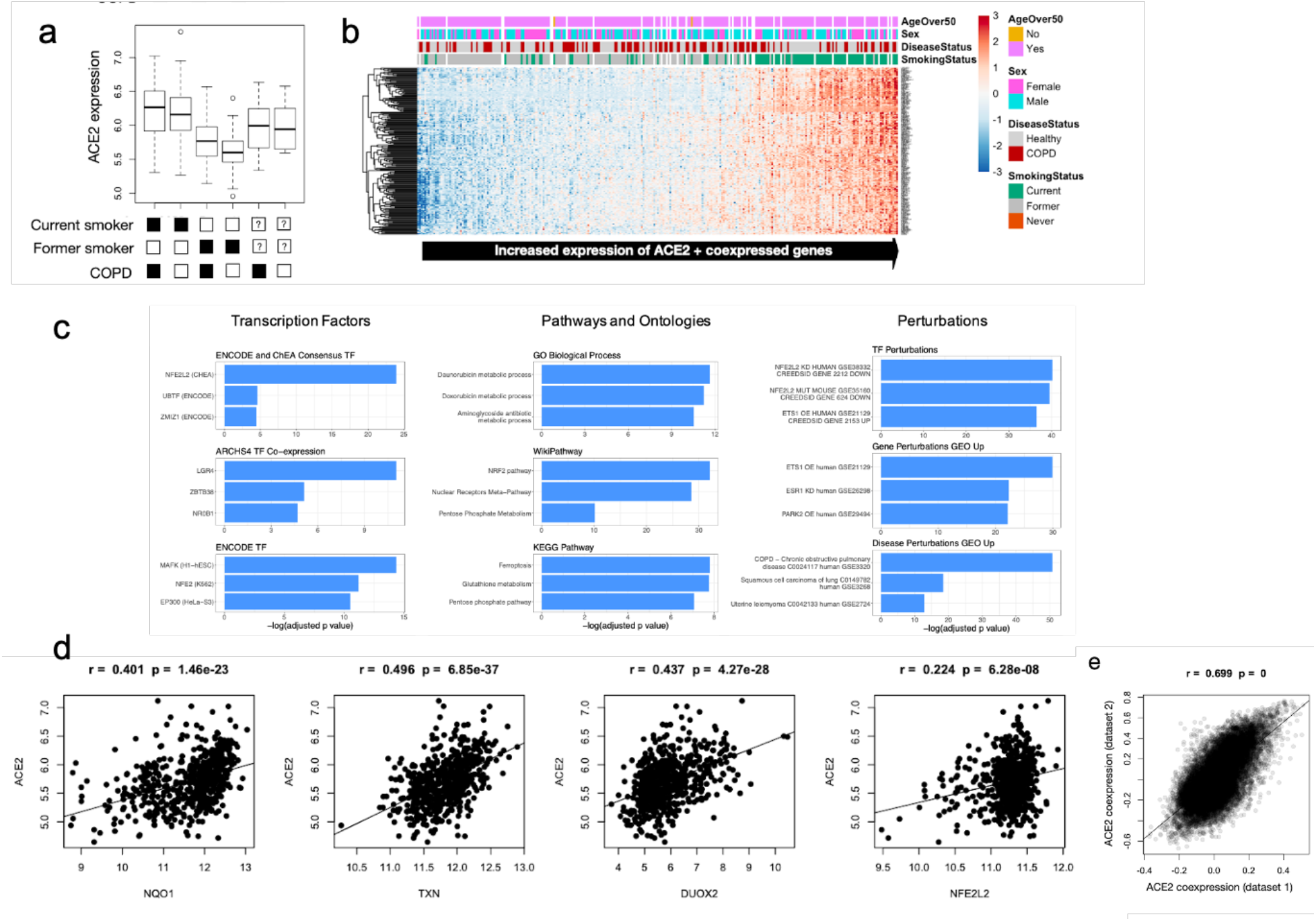
Expression and functional enrichment analysis of *ACE2* and co-expressed genes in smokers and individuals with COPD. **a)** Analysis of relative *ACE2* expression with respect to smoking status and COPD diagnosis. **b)** Expression of top 200 *ACE2*-correlated genes (including *ACE2*) individuals with various smoking status and COPD diagnosis (N=345). **c)** Functional enrichment analysis of top 200 *ACE2*-correlated genes (including *ACE2*). Terms are ranked by -log_2_(FDR-adjusted *p* value) for nine ontologies/groups of interest. **d)** Pearson correlation of *ACE2* with important interferon-related candidate genes found to be co-expressed with *ACE2*. **e)** Correlation between dataset 1 (N=159, Figure 3) and dataset 2 (N=345, Figure S3).

**Figure S4.**
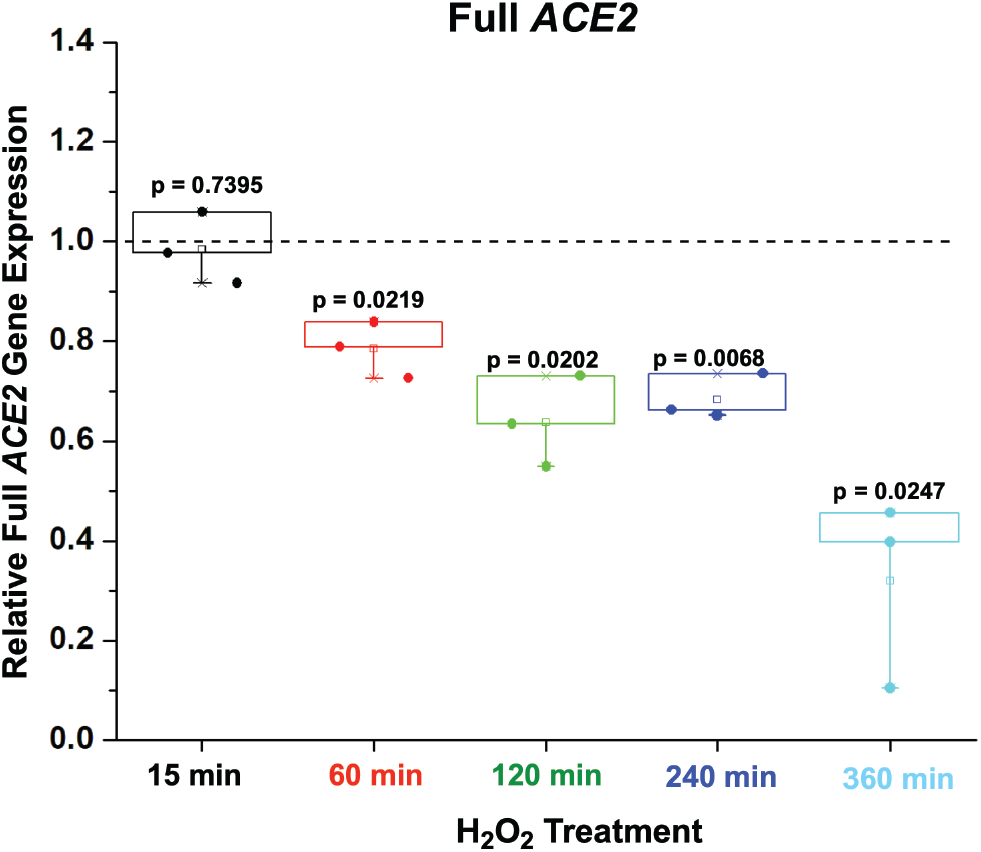
*ACE2* expression in response to oxidative stress. Expression of *ACE2* in Calu-3 cells (relative to untreated controls) following H_2_O_2_ treatment for the indicated length of time (see Methods, Table S2).

